# Dengue Disease Dynamics are Modulated by the Combined Influence of Precipitation and Landscapes: A Machine Learning-based Approach

**DOI:** 10.1101/2020.09.01.278713

**Authors:** Micanaldo Ernesto Francisco, Thaddeus M. Carvajal, Masahiro Ryo, Kei Nukazawa, Divina M. Amalin, Kozo Watanabe

## Abstract

**Background:** Dengue is an endemic vector-borne disease influenced by environmental factors such as landscape and climate. Previous studies separately assessed the effects of landscape and climate factors on mosquito occurrence and dengue incidence. However, both factors interact in time and space to affect mosquito development and dengue disease transmission. For example, eggs laid in a suitable environment can hatch after being submerged in rain or flood water.

**Objectives:** This study aimed to investigate the combined influences of landscape and climate factors on mosquito occurrence and dengue incidence.

**Methods:** Entomological, epidemiological, and landscape data from the rainy season (July-December) were obtained from respective government agencies in Metro Manila, Philippines, from 2012 to 2014. Temperature, precipitation, and vegetation data were obtained through remote sensing. A random forest algorithm was used to select the landscape and climate variables. Afterwards, using the identified key variables, a model-based (MOB) recursive partitioning was implemented to test the combinatory influences of landscape and climate factors on the ovitrap index and dengue incidence.

**Results:** The MOB recursive partitioning for the ovitrap index indicated that mosquito occurrence was higher in high residential density areas, where industrial areas also exist and are well connected with roads. Precipitation was another key covariate modulating the effects of landscape factors, possibly by expanding breeding sites and activating mosquito reproduction. Moreover, the MOB recursive partitioning indicated that precipitation was the main predictor of dengue incidence, with a stronger effect in high residential density and commercial areas.

**Discussion:** Precipitation with floods has epidemiologically important implications by damaging shelters and causing population displacement, thus increasing exposure to dengue vectors. Our findings suggest that the intensification of vector control during the rainy season can be prioritized in residential and commercial areas to better control dengue disease dynamics.

## INTRODUCTION

Dengue is an endemic vector-borne disease influenced by environmental factors such as climate and landscape. Dengue-endemic countries such as the Philippines consider this arboviral disease an economic and health burden (Buczak, et al. 2014). Environmental factors, particularly climate and landscape, play a significant role in regulating the temporal variation and spatial distribution of dengue and the vectors *Aedes aegypti and Ae. albopictus* (Hayden, et al. 2010). These factors can mediate human-mosquito interactions by expanding the vector’s habitat and increasing its abundance, thus advancing dengue disease transmission (Thongsripong, et al. 2013).

Previous studies demonstrated that climate factors such as precipitation and temperature significantly affect both mosquito abundance (Barrera, Amador and MacKay 2011, Naish, et al. 2014) and dengue incidence (Phanitchat, et al. 2019, Carvajal, Viacrusis, et al. 2018). For example, the high availability of breeding sites for mosquitoes during the rainy season in Southeast Asian countries (e.g., Philippines, Singapore, Thailand, and Indonesia) contributes to the increased number of annual dengue cases (Su 2008, Hashizume, et al. 2012). Many studies have reported that the increasing number of cases is associated with the high number of available mosquito breeding sites that can hold or contain rainwater, thereby facilitating high mosquito abundance (Seidahmed, et al. 2018, Arcari, Tapper and Pfueller 2007). Additionally, high temperatures are responsible for extending adult mosquito longevity, accelerating virus replication, and enhancing the mosquito biting rate (Kilpatrick, et al. 2008, Chan and Johansson 2012).

Recent studies have shown that different land use (LU) types (residential, industrial, and agricultural areas) may have different impacts on dengue incidence (Kesetyaningsih, et al. 2018, Sheela, et al. 2017, Sarfraz, et al. 2012, Vanwambeke, Lambin, et al. 2007, Cheong, Leitão and Lakes 2014) given the uneven spatial distribution of vectors in these terrestrial habitats (Piovezan, et al. 2019). Areas with human settlements contribute to a high incidence of dengue (Cheong, Leitão and Lakes 2014, Sarfraz, et al. 2012) due to the high availability of man-made water-holding containers that serve as breeding sites (Ngugi, et al. 2017) and humans as a host preference for blood meals (Higa 2011).

Most previous studies investigated the effects of either dynamic climate factors (Carvajal, Viacrusis, et al. 2018, Zheng, et al. 2019, Arcari, Tapper and Pfueller 2007, Tovar-Zamora, et al. 2019, Bavia, et al. 2020) or static spatial distributions of landscape attributes (Seidahmed, et al. 2018, Vanwambeke, Lambin, et al. 2007, Vanwambeke, N.Bennett and Kapan 2011, Sarfraz, et al. 2012) on the temporal variations or spatial distributions of mosquito occurrence and dengue incidence. However, landscape and climate conditions can change concurrently in time and space, and their spatiotemporal interrelation may play a decisive role in determining the positive or negative effects on mosquitoes and dengue, such as interference during flood events. In small areas where rainfall is equally distributed, rainwaters flow from highlands to lowlands due to gravity, and patterns of water flow and retention can greatly differ in LU type due to surface roughness texture and permeability. Comparative studies reported high mosquito densities in flooded lowlands compared with non-flooded highlands (Nasir, et al. 2017, Rydzanicz, Kącki and Jawień 2011). One study reported that the high mosquito abundance in lowlands was influenced by floods that reach mosquito eggs that were previously laid in the environment (Hashizume, et al. 2012). Another study demonstrated that during the dry season, mosquito abundance was high in residential areas given the availability of permanent water-holding containers that served as breeding sites (Little, Bajwa and Shaman 2017); in the wet season, mosquito reproduction expanded to other non-residential areas. These studies found an uneven effect of precipitation on mosquito abundance due to different ecological responses in different local landscapes (Nasir, et al. 2017, Rydzanicz, Kącki and Jawień 2011, Little, Bajwa and Shaman 2017). The characteristics of a local area’s landscape can also influence its microclimate (Chang, Li and Chang 2007, Lin, et al. 2018, Thani, Mohamad and Abdullah 2017, Shashua-Bar, Pearlmutter and Erell 2011), potentially thus affecting the ecology of the mosquito and dengue transmission. For example, areas with a high percentage of impervious surfaces (e.g., paved roads, built-up areas) with less vegetation coverage can absorb high amounts of solar radiation and produce more heat compared to areas with less impervious surfaces and extensive vegetation coverage (Koch-Nielsen 1999). Therefore, the combinatory influence of landscape and climate factors on mosquito and dengue incidence must be quantitatively assessed (Sallam, et al. 2017). No studies have yet attempted to assess the combined influence of climate and landscape features on dengue disease dynamics.

Previous studies that utilized environmental factors to develop dengue epidemiology models faced challenges when jointly considering climate and landscape attributes, preventing us from better understanding dengue disease distribution. One such challenge is the availability of secondary datasets (Sarfraz, et al. 2012, Vanwambeke, Lambin, et al. 2007). Climate data such as temperature and precipitation are typically obtained from ground weather stations. However, using such data is limited by the limited number of ground weather stations. Therefore, remotely sensed climatic variable data have been utilized in epidemiological studies to address the lack of routinely collected data from ground meteorological stations (Kapwata and Gebreslasie 2016, German, et al. 2018). With the introduction of platforms that integrate remote sensing and cloud computation such as Google Earth Engine (GEE) (Gorelick, et al. 2017), it is possible to freely access and process many types of satellite-derived products with notable flexibility, even in large areas (DeVries, et al. 2020). However, many studies that utilize LU based on satellite images contain certain limitations. In this type of map, built-up areas are often treated as a single category (Vanwambeke, et al. 2006, Ibarra, et al. 2014, German, et al. 2018), preventing the ability to further distinguish the subcategories of land utilization such as residential, commercial, industrial, etc. These different categories of LU have different ecological responses to mosquito and dengue dynamics (Thammapalo, et al. 2007); hence, detailed maps might amplify the chances to capture fine scale variations of mosquito habitats and dengue incidence. Although labor intensive, detailed LU maps produced by local governmental agencies based on field observations can help uncover patterns of dengue disease at a fine scale by capturing the different categories of land utilization in urban areas (Nazri, et al. 2011).

Another challenge is finding an appropriate method to model complex interactive mechanisms between multiple environmental factors (Little, Bajwa and Shaman 2017, Sarfraz, et al. 2012). In the recent decade, modeling techniques in machine learning methods such as random forests (RFs) (Breiman 2001) have been adopted to analyze complex databases and handle anomalies found in datasets such as outliers and multi-collinearity among covariates. Data-intensive modeling has gained popularity in spatiotemporal ecological modeling at the landscape or larger scales to better explain ecological or epidemiological patterns by capturing nonlinear variable interactions (Ryo, Yoshimura and Iwasaki 2018, Ryo, Harvey, et al. 2017b, Ryo and Rilling 2017a). The results of this approach improved RF model accuracy (Leontjeva and Kuzovkin 2016) and better predictability of species’ habitat distribution with the inclusion of maximum entropy (Stanton, et al. 2012).

This study aimed to examine the combinatory influence of landscape and climate features on mosquito occurrence and dengue incidence across Metropolitan Manila, the Philippines. We employed some advanced machine learning algorithms due to its growing utilization to explore the influence of landscape features or climate on dengue disease (Carvajal, Viacrusis, et al. 2018, Guo, et al. 2017, Ong, et al. 2017, Chen, et al. 2018, Baquero, Santana and Chiaravalloti-Neto 2018) and mosquito occurrence (Mwanga, et al. 2019, Jiménez, et al. 2019, Früh, et al. 2018, Zheng, et al. 2019). By selecting important environmental features for RFs, we further examined and described the optimal combination of landscape and climate conditions that influence dengue disease and mosquito occurrence using model-based (MOB) recursive partitioning.

## MATERIALS AND METHODS

### Study area

Metropolitan Manila is the National Capital Region (NCR) of the Philippines, located at Southwestern Luzon (14°50’N Latitude, 121°E Longitude). With 100% urbanization (Asian Development Bank 2014), the NCR is the most densely populated area in the country (18,165.1 persons/km^2^ spread over an administrative land area of 636 km^2^) (Asian Green City Index 2011). It comprises 16 cities and one municipality with a total population of 12,877,253 (Philippines Statistics Authority 2019). Each city or municipality is further subdivided into the smallest administrative division, a “Barangay,” commonly known as a village, with 1,706 total villages. A collection of villages can be merged into a “zone” depending on the city’s administrative boundaries.

The majority of the target area is covered by residential (54.07%), industrial (9.41%), and commercial (7.45%) areas. The territorial development of Metro Manila occurred through a gradual replacement of agricultural LU by industrial, commercial, and a massive increase in residential areas. The constant spatial and population growth has led to LU pressure and instigated substandard housing in areas with a high risk of flooding (Zoleta-Nantes 200).

The climate in Metro Manila is divided into two major seasons: the dry season (November to April) and the wet season (the rest of the year) (DOST 2014). The rainy season from June to September is characterized by strong monsoon rain and tropical storms (World Bank 2014). Heavy rain associated with a lack of drainage infrastructure leads to a high vulnerability for flooding (Zoleta-Nantes 200).

### Data Sources and Processing

#### Administrative boundaries

The map of the administrative boundaries of Metropolitan Manila (Figure 1a) was obtained from the Philippine GIS Data clearinghouse (www.philgis.org). Metropolitan Manila includes 1,706 villages (barangays) with most within the City of Manila (n = 897; 53%). In this study, the villages of Manila, Caloocan, and Pasay were merged together into “zones” to facilitate consistency in village size because most villages are very small with an average area of 0.06 km. Additionally, 86% (n = 771) of the villages have a total area of <0.06 km^2^. The average total area of each village in Metropolitan Manila (excluding the City of Manila) is 0.41 km^2^. This study used the City of Manila, Caloocan, and Pasay’s designated zone names to merge villages. Overall, 464 villages or zones were subsequently analyzed in this study. The population statistics were obtained from the Philippine Statistics Authority agency (www.psa.gov.ph). Since the Philippine population census is conducted every five years, we obtained the 2010 (Philippines Statistics Authority 2012) and 2015 (Philippines Statistics Authority 2019) census data and used the compounded population growth rate to calculate the population for the years 2012 and 2013. The sum of the projected population of the merged villages (Manila, Caloocan, and Pasay) was also calculated.

**Figure 1:**
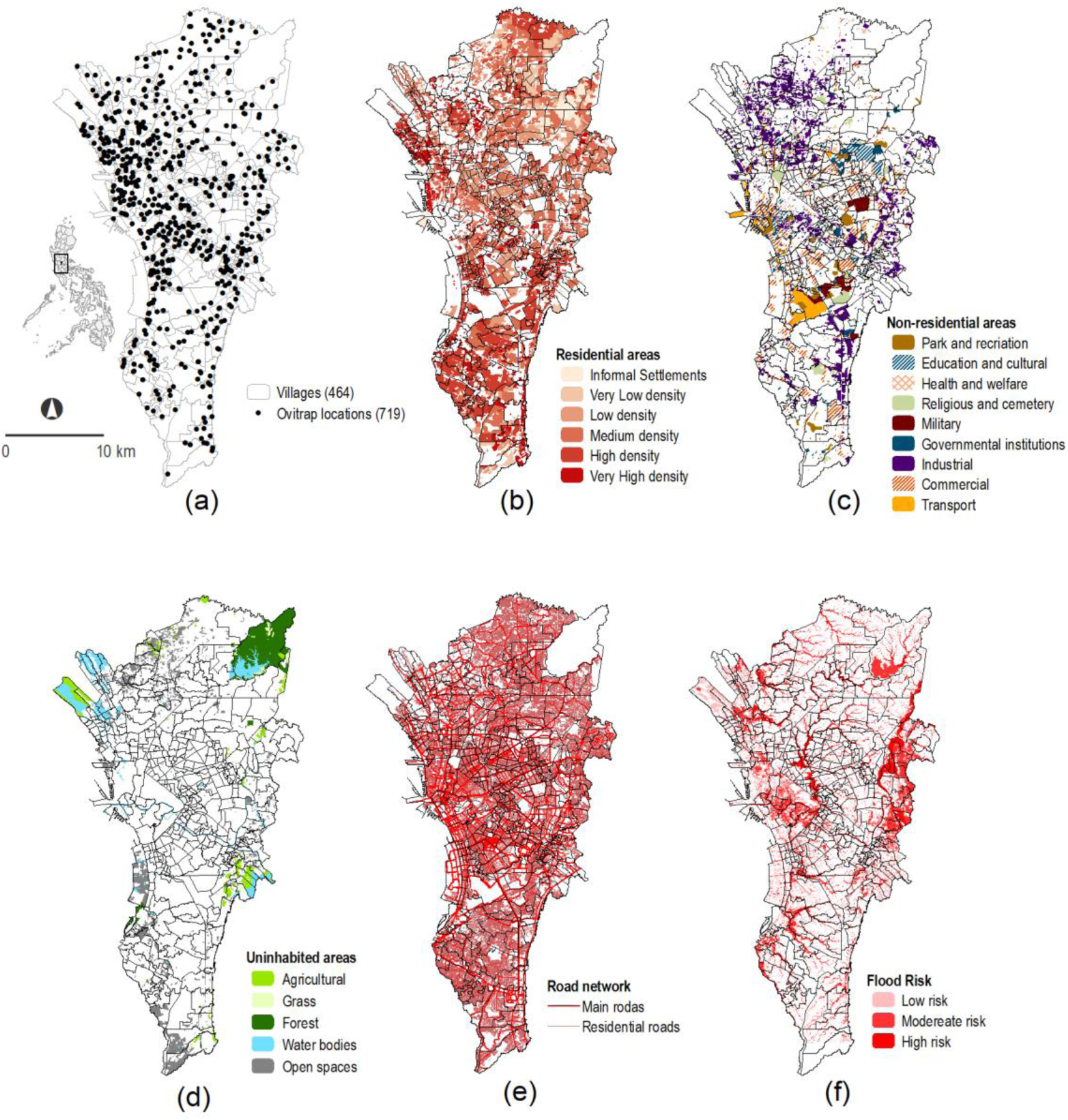
Administrative boundaries of Metropolitan Manila showing: (a) Ovitrap locations; Landscape features: (b) Residential areas classified according to densities, (c) Non-residential areas, (d) Uninhabited areas, (e) Road networks, and (f) Flood risk.

#### Entomological surveillance

In 2012, governmental institutions (Department of Science and Technology (DOST), Department of Education, Department of Health, Department of Interior, and local governments) implemented a nationwide surveillance program that installed DOST Orvicidal/Larvicidal traps (OL-traps) to monitor *Aedes* mosquitoes to help control dengue transmission and reduce dengue cases (DOST n.d., DOST Mosquito Ovicidal/Larvicidal (OL) Trap for Dengue Prevention 2013). Surveillance programs in many countries have utilized ovitraps as a routine surveillance tool because they are relatively low-cost and reliable in attracting gravid *Aedes* females for oviposition (Silver 2007, Ritchie, et al. 2003). In Metro Manila, ovitraps were installed in public places such as schools, institutes, and other education facilities. A total of 719 georeferenced surveillance locations containing reported Ovitrap indices (OIs) were extracted from the reporting website (http://oltrap.pchrd.dost.gov.ph/) (DOST n.d.) from July 2012 to December 2014. Afterwards, each georeferenced surveillance location was matched with its corresponding village, with only 268 of the 464 villages containing mosquito surveillance location(s). We aggregated the OIs into a monthly index by dividing the cumulative OI by the total number of sampling locations. Given the low numbers and inconsistent reporting during the months of January to June, the study only included the aggregated monthly OI from July to December of 2012, 2013, and 2014 (Additional file 3).

#### Epidemiological data

The total number of weekly reported dengue cases from January 2012 to December 2014 for all 464 villages was obtained from the National Epidemiology Center, Department of Health, Philippines. We calculated the monthly dengue incidence by dividing the total number of dengue cases each month by the total population of the village multiplied by a population factor of 10,000. The dengue incidence was transformed by adding 1 to all values and obtaining its natural logarithm [loge (n + 1)].

#### Climatic factors

Remote sensing (RS) is a promising tool in epidemiological studies (German, et al. 2018, Misslin and Daudé 2017, Buczak, et al. 2014, Araujo, et al. 2015). This study used the Tropical Rainfall Measurement Mission (TRMM) product 3B43 to obtain the monthly average rainfall. This gridded quasi-global product consists of monthly average precipitation measured in hourly bases with 0.25° of spatial resolution (Huffman and Bolvin 2018). The Terra Moderate Resolution Image Spectroradiometer (MODIS) collected the average land surface temperature. The product MOD11A2 consists on the average temperature collected within an eight-day period for both daytime and nighttime temperatures with 1 km spatial resolution (USGS n.d.). GEE (Gorelick, et al. 2017) was used to download the RS raster images, apply scaling factor (0.02), and convert temperature values from the default Kelvin (K) to degrees Celsius (°C). This product suffers from missing data, particularly during the rainy season (August-October), given the high cloud cover and other atmospheric disturbances. To overcome this limitation, a Kriging interpolation method was applied to complete monthly temperature records for each village using ArcGIS software version 10.2 (ESRI, Redlands, CA). This method weights the surrounding measured values to derive a surface of predicted values for an unmeasured location (ESRI 2016). Since each village can be covered by multiple pixels of the raster images of precipitation and temperature, the mean value of all pixels within each village (spatial) was calculated per month (temporal) and used for further analyses.

A flood hazard map of Metro Manila was obtained from the LiDAR Portal for Archiving and Distribution (LiPAD) website (https://lipad.dream.upd.edu.ph) (LiPAD 2018). This flood map indicates the flood susceptibility level at a 10-m spatial resolution (NOAH 2015). There are three categories of flood susceptibility: (a) low (flood water height ranging from 0.1–0.5 m), b) moderate (0.5–1.5 m), and (c) high (above 1.5 m). Initially, the percentage of land covered by each flood risk category was calculated by village and multiplied by a weighing value from 1 (low) to 3 (high) according to the risk category. The average of these three values was calculated and utilized as the flood risk index per village. The degree of flood susceptibility was estimated based on a five-year period of heavy rain scenarios and thus is limited to spatial risk and does not consider the variation of risk throughout a year.

#### Landscape factors

The local LU map of Metropolitan Manila (2004) was obtained from the Philippine Geoportal website (www.geoportal.gov.ph) managed by the Bureau of National Mapping and Resource Authority and the Metropolitan Manila Development Authority (NAMRIA n.d.). This map contains 30 LU types (agricultural, grass, forest, water bodies, open spaces, parks and recreation, education and cultural, health and welfare, religious and cemetery, military, governmental institutions, industrial, commercial, transport, residential areas of very low, low, medium, high, and very high density, and informal settlements). The largest portion (54.07%) was covered by residential areas (very low, low, medium, high, and very high densities) with multi-story dwelling places (1–2, 3–4, 5, or more stories). This study only considered the house density categories (very low, low, medium, high, and very high) given the very small proportion of multi-story categories of more than three stories (0.01%–0.07%). Non-residential areas such as industrial, commercial, and public facilities comprised 37% of Metropolitan Manila. A small portion was covered by natural landscape aspects such as water bodies and forests (10%). The 2004 LU map had a dengue incidence time gap (2012–2014); thus, we updated the map to the period covered by our study so that all input parameters in the model had the same time range. This map was subjected to updates based on open street maps (OSM), processed and distributed by Geofabrik GmbH (www.geofabrik.de). The OSM data contained modifications that occurred before December 2016 (Geofabrik GmbH 2019). Prior knowledge of the study area was used to manually inspect and validate the map modifications. LU variables included the percentage of land covered by each LU class (i.e., agriculture, water bodies, commercial, residential) per village (Figure 1b–d). The percentage of each LU class was calculated as follows. Firstly, we calculated the area of each LU class per village. Then, the percentage of each class was determine over the total area of the villages. All edits and calculations of LU areas per village was performed in ArcGIS software, version 10.2.

Road network density (RND) assesses the urbanization gradient (Suarez-Rubio and Krenn 2018), which influences mosquito abundance and dengue transmission (Bostan, et al. 2017). The road network map was obtained from the Philippine GIS Data Clearinghouse website (http://philgis.org/) and classifies roads as primary, secondary, tertiary, residential, and others (PhilGIS 2012). The RND for each category was calculated by dividing the total length of roads by the total village area. Since the RND of each category of primary, secondary, and tertiary roads was less than 0.001 m/m^2^, we merged them into a single category, “main roads” (Figure 1e). Terra MODIS Normalized Difference of Vegetation Index (NDVI) was derived from the product MOD13Q1 version 6. The NDVI consists of measures of the reflected photosynthetic activity on vegetation and is generated every 16 days at 250-m spatial resolution (USGS n.d.). All images were downloaded through GEE and processed using ArcGIS to obtain their monthly averages per village.

#### Cross-correlation analysis

A cross-correlation analysis was conducted on the temporal variations of environmental factors (precipitation, temperature, and vegetation) on the OI and dengue incidence. The mean value of Metropolitan Manila area per month for each variable was utilized. We identified the best-lag based on the highest Pearson correlation coefficient that was generated and its statistical significance (*p* < 0.05). These analyses were implemented in R software version 3.6.2 using “*ggpubr”* package version 0.2.4 (Kassambara 2019). The best-lag timing for each variable was used for the latter analyses.

#### Model development with variable selection

RF is a bootstrap aggregation (bagging) ensemble method that generates a large number of independent bootstrapped trees from random small subsets of the dataset (Breiman 2001). RF is used to solve a variety of classification and regression problems due its ability to handle large numbers of predictor variables even in the presence of complex interactions (Garge, Bobashev and Eggleston 2013). Two regression models were implemented in this study. Dengue incidence was regressed with lagged climate factors, LU types, and OI, with 27 explanatory variables. Additionally, the OI was regressed with lagged climate factors and LU types, with 26 explanatory variables. RF models were estimated using the “*ranger*” package (Wright, Wager and Probst 2020) implemented in the R software version 3.6.2 (R Core Team 2017) with parameters set at *ntree* = 500 and *mtry* = 5.

To identify the most important predictors of dengue incidence and mosquito occurrence, we assessed variable importance (VI), which was measured as the mean decrease in MSE in the RF models. VI is calculated based on the number of times the explanatory variable is used for splitting, weighted by the improvement to the model as a result of each split, averaged over all trees (Elith, Leathwick and T.Hastie 2008). For the VI and respective p-values, we applied the permutation importance method, which computes an unbiased VI measure (Zeileis, Hothorn and Hornik 2008). Positive importance values with p-values less than 0.05 were selected for the subsequent MOB recursive partitioning analysis.

To investigate the interactions of the selected explanatory variables, we used MOB recursive partitioning to build a decision tree hierarchy by recursively partitioning the data into several groups using the variable that provides the best split per node (Zeileis, Hothorn and Hornik 2008). Recursive partitioning was implemented via two steps. First, we utilized all selected explanatory variables to build a decision tree to see the overall distributions of the OI and dengue incidence. From this step, we can also identify the main predictors of the OI and dengue incidence. In the second step, we explicitly specified the most important predictor found in the earlier step as the main explanatory variable in the model and then explored covariates that modulate the associations between the main predictors and the response variables. Herewith, we inferred how the OI and dengue incidence were modulated by the interaction between the main predictor and other variables. We used the Linear Model Tree (*lmtree*) interface implemented in “*partykit*” package (Hothorn and Zeileis 2015) in R Software (R Core Team 2017). For each of the generated intermediate nodes, regression coefficients and *p-values* were computed to inform the relevance of the splitting variable in a particular node. We set *p value* < 0.05 and maximum depth = 4 as the stopping criteria so that only the most important variables and intermediate nodes that were statistically significant were used for modeling. These hyperparameters have been suggested to avoid model overfitting (Zeileis, Hothorn and Hornik 2008, Pirkle, et al. 2018), and simplification of tree structure was performing by only including the most relevant predictors (Kopf, Augustin and Strobl 2010). The *p-values* were *Bonferroni* corrected to control a false positive rate. The slope coefficients, R-squared, and p-values were obtained using the *“summary()”* function.

## RESULTS

### Cross-correlation analysis

Precipitation yield had the highest positive and significant correlation with dengue incidence (r = 0.69, p = 0.00) at a one-month lag, followed by OI (r = 0.52, p = 0.05) and temperature (r = 0.52, p = 0.05), both at a three-month lag. Vegetation displayed a negative and significant correlation (r = −0.71, p = 0.00) at a one-month lag (Table 1). Vegetation showed the highest positive correlation with the OI (r = 0.77, p = 0.00) at a three-month lag, followed by temperature (r = 0.73, p = 0.00) and precipitation (r = 0.48, p = 0.04), both at a zero-month lag.

**Table 1:** Cross-Correlation Analysis of Temporal Climate Factors in Dengue Incidence and Ovitrap Index

| Variables | Dengue incidence |  |  | Ovitrap index |  |  |
| --- | --- | --- | --- | --- | --- | --- |
|  | *Lag month* | *r value* | *p value* | *Lag month* | *r value* | *p value* |
| Ovitrap Index | 3 | 0.52 | $p = 0.05$ | - | - | - |
| Precipitation | 1 | 0.69 | $p = 0.00$ | 0 | 0.48 | $p = 0.04$ |
| Temperature | 3 | 0.52 | $p = 0.05$ | 0 | 0.73 | $p = 0.00$ |
| Vegetation | 1 | -0.71 | $p = 0.00$ | 3 | 0.77 | $p = 0.00$ |

### Variable selection

Figure 2 shows the variable importance of the selected variables in the two RF models. The OI was significantly associated with 11 variables (three climatic factors and eight landscape factors). High residential density areas were ranked first, followed by precipitation, residential RND, temperature, medium residential density areas, vegetation, health institution areas, very high residential density areas, flood risk, commercial areas, and industrial areas (Figure 2a). Dengue incidence was significantly associated with eight variables (three climatic factors, four landscape factors, and OI). Precipitation was ranked first, followed by temperature, commercial areas, high residential density areas, vegetation, OI, flood risk, and residential RND (Figure 2b). These variables were used in the subsequent modeling.

**Figure 2:**
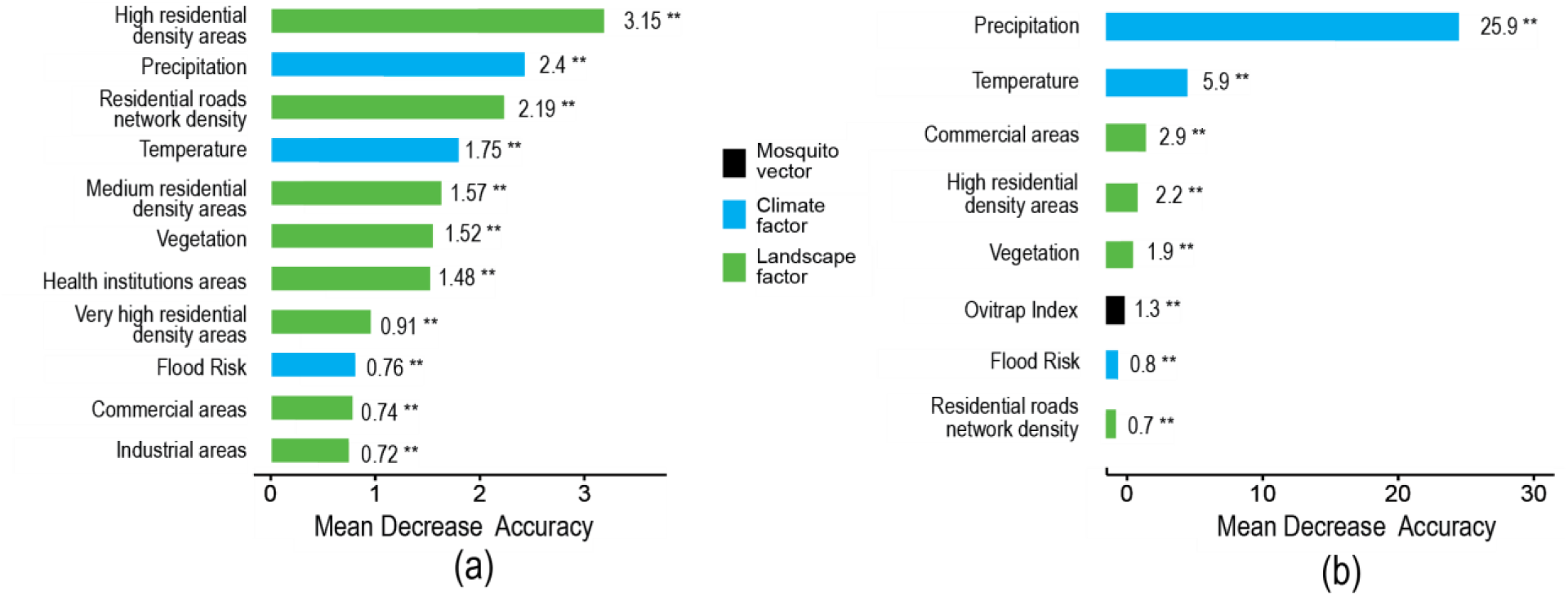
Variable importance measures of the variables with the most significant associations with (a) ovitrap index and (b) dengue incidence; (**) statistically significant at p < 0.05.

### Model-based recursive partitioning

Figure 3a displays the MOB tree of the environmental conditions explaining the distribution of the OI. The tree is composed of three partitioning levels and eight terminal nodes. Each terminal node shows the average OI of a subset of the entire dataset based on selected landscape and climatic features and labeled accordingly as terminal nodes OV-A1 to OV-A8. In the MOB tree, high residential density was identified as the first-level partitioning variable and thus considered the most important environmental feature. The succeeding levels were comprised of residential RND, precipitation, industrial areas, health institutional areas, and flood prone areas. The order of the variables was agreed with the estimated variable importance (Figure 2a). The average OI from node OV-A1 to OV-A8 ranged from 1.36 to 34.52%.

**Figure 3:**
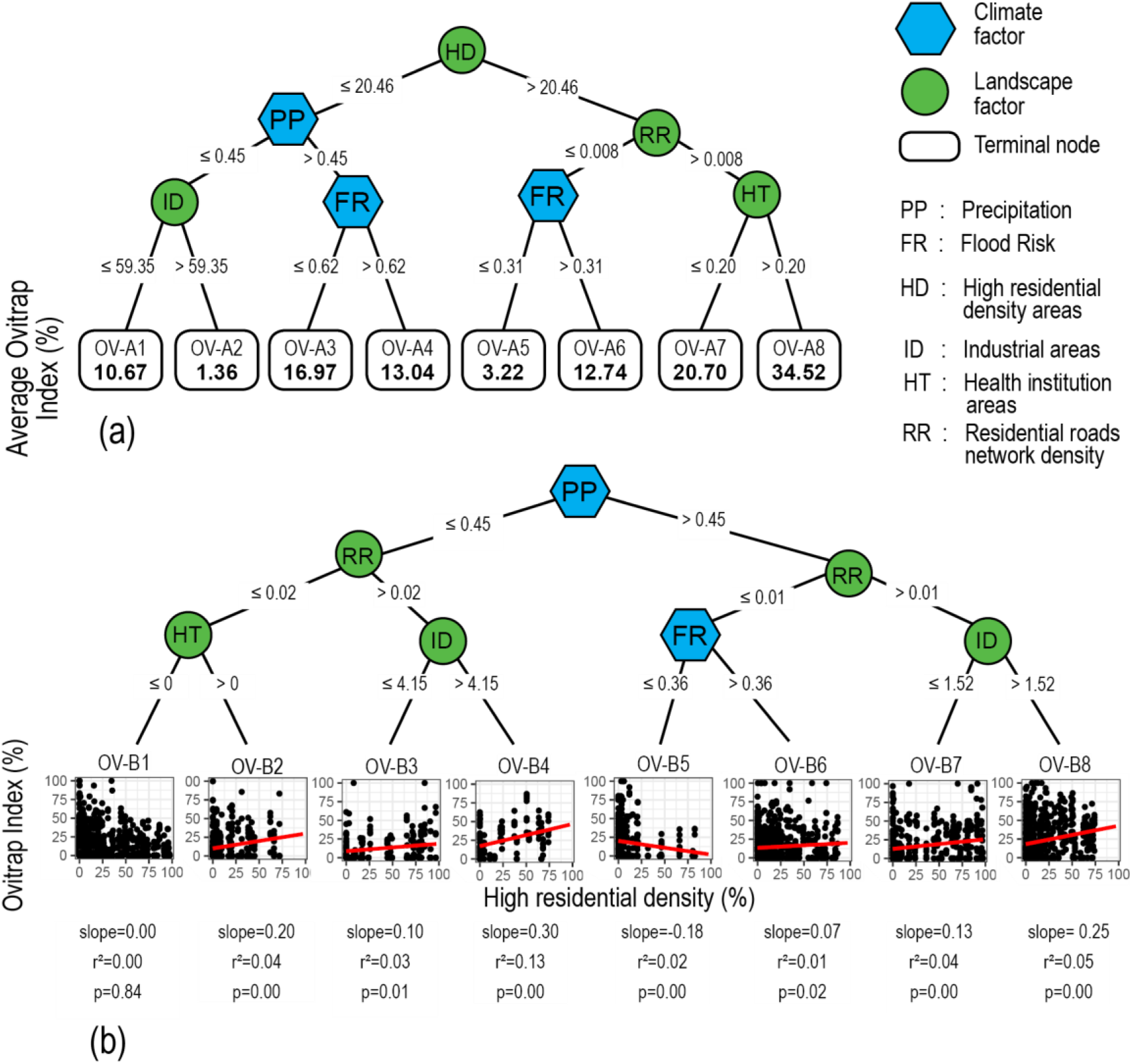
Recursive partitioning trees for identifying the (a) most influential variables on the ovitrap index and (b) interactive effects between environmental factors and the ovitrap index.

We employed further analyses to identify the interactive effects of the most important predictor (i.e., high residential density) with other environmental factors on OI (Figure 3b). Higher slopes and R-squared values were found in nodes OV-B4 and OV-B8 (0.30 (R^2^ = 0.13, p = 0.00) and 0.25 (R^2^ = 0.05, p = 0.00), respectively), indicating that the effect of high residential density areas on OI is modulated by precipitation, residential roads, and industrial areas. Discordant associations of high residential density areas to OI were observed for flood risk. A negative slope of −0.18 (R^2^ = 0.02, p = 0.00) was found when the flood risk was lower or equal to 0.36 (node OV-B5) whereas a positive slope of 0.07 (R^2^ = 0.01, p = 0.02) was found when the flood risk was greater than 0.36 (node OV-B6).

Figure 4a shows the MOB tree of the influence of climatic and landscape factors on dengue incidence. This tree is composed of eight terminal nodes generated from three partitioning levels. Each terminal node shows the average dengue incidence of a subset of the entire dataset based on selected climatic and landscape factors and is labeled accordingly as terminal nodes DI-A1 to DI-A8. Precipitation was the partitioning variable in the first and second partitioning levels and thus considered the most important environmental feature, similar to the RF analysis (Figure 2b). The succeeding partitioning levels comprised commercial and high residential density areas. The average dengue incidence on the terminal nodes DI-A1 to DI-A8 ranged from 0.04 to 0.51.

**Figure 4:**
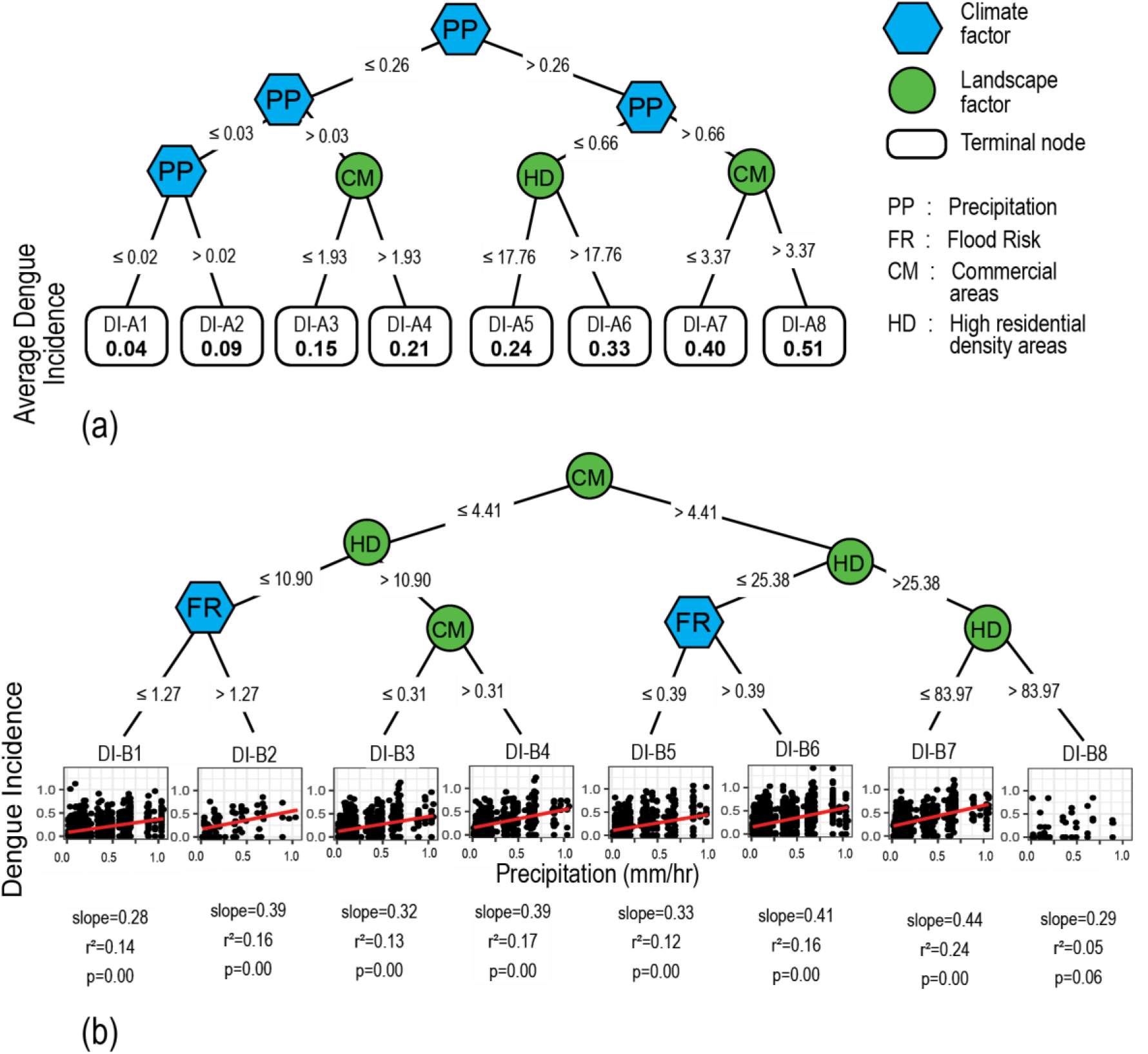
Recursive partitioning trees for identifying the (a) most influential variables on dengue incidence and (b) the interactive effects between environmental factors and dengue incidence.

We employed further analyses to infer the interactive effect of the most important predictor (precipitation) with other environmental factors on dengue incidence (Figure 4b). We specified precipitation as the main predictor and the remaining variables as interacting factors. Higher slopes and R-squared values were found in relation to two different interaction patterns and were considered influential toward dengue incidence. The first pattern involved interactions between precipitation, commercial areas, and high residential density areas (nodes DI-B7 and DI-B4, with slopes of 0.44 (R^2^ = 0.24, p = 0.00) and 0.39 (R^2^ = 0.17, p = 0.00), respectively). The second pattern involved interactions between precipitation, commercial areas, high residential density areas, and flood risk (nodes DI-B6 and DI-B2, with slopes of 0.41 (R^2^ = 0.16, p = 0.00) and 0.39 (R^2^ = 0.16, p = 0.00), respectively).

## DISCUSSION

### The interactive effects between high residential density, precipitation, and other landscapes in modulating ovitrap index

Ovitraps, in general, can detect the presence of both Ae. aegypti and Ae. albopictus. However, the OI data utilized in this study does not contain any information on the proportion of these two species. Therefore, in this section, our discussion focuses solely on *Ae. aegypti*. This is because previous studies that surveyed selected areas of Metropolitan Manila indicated a high infestation rate of *Ae. aegypti* (>80%) (Mistica, et al. 2019, Carvajal, Ho, et al. 2019), thereby making it the primary vector for dengue transmission.

Both the RF and MOB tree (Figures 2a and 3a) analyses clearly indicate the importance of high residential density areas on the overall distribution of the OI. This finding is consistent with previous studies reporting that high residential areas are potential sources of breeding sites and blood meals for *Ae. aegypti* (Ngugi, et al. 2017). These areas are highlighted as the foremost breeding sites for *Ae. aegypti* due to the permanent availability of breeding containers and humans, who serve as blood sources (Higa 2011, Ngugi, et al. 2017, Little, Bajwa and Shaman 2017). Herewith, residential areas exhibited ecological requirements to serve as main mosquito habitats compared with other types of landscapes.

To determine how high residential density areas interact with other environmental factors in modulating OI, we employed further analyses by specifying high residential density areas as the main predictor and the remaining variables as interacting factors (Figure 3b). The influence of high residential density areas on mosquito occurrence became clearer under certain environmental conditions. The two nodes OV-B4 and OV-B8 with the highest slopes (0.30 and 0.25, respectively) were formed in high industrial areas (>4.15 and >1.52%, respectively) and high residential road areas (>0.02 and >0.01 m/m^2^, respectively), and node OV-B4 had a low precipitation condition (≤0.45 mm/hr). Since *Ae. aegypti* preferentially breeds in small water containers exposed to the outdoors (Carvajal, Ho, et al. 2019, Ngugi, et al. 2017), little rainfall might be sufficient to maintain optimal levels of water suitable for mosquito emergence. Conversely, enhanced rainfall might flush out mosquito eggs and larvae from breeding containers and reduce the sensitivity of the OI to changes in high residential density areas. The destructive effect of significant precipitation on mosquito development has been suggested in another work (Dickin, Schuster-Wallace and Elliott 2013); however, the possible precipitation thresholds remain unclear.

Residential RND was the partitioning variable on the second level of the MOB tree. Furthermore, nodes OV-B4 and OV-B8, which had the highest slopes, were partitioned with higher residential road densities (>0.02 and >0.01 m/m^2^, respectively). These results suggest an enhanced vulnerability to mosquito occurrence in high residential density areas with a higher density of residential roads. Roads not only serve as transportation networks for people and goods but are also simultaneously accompanied by drainage components (e.g., roadside drains or canals, drain sumps), which collect surface water runoff for discharge in appropriate locations to avoid flooding. However, in many cases, efficient drainage in residential areas can be compromised by the encroachment of concrete structures or garbage clogging canals (Lagmay, et al. 2015). These interferences can inhibit complete water flow, resulting in spots of accumulated water, which can create favorable habitats for *Ae. aegypti* (Paploski, et al. 2016). Growing evidence has suggested a positive association between drainage and the occurrence of *Ae. aegypti* in Singapore (Seidahmed, et al. 2018), Brazil (Souza, et al. 2017)m and Australia (Montgomery, et al. 2004).

Notably, with increased precipitation (>0.45 mm/hr), high residential density areas showed an opposite association to OI depending on the flood risk (Figure 3b). Increased flood risk led to a positive association between high residential density areas and OI (node OV-B6) whereas reduced flood risk led to a negative association (node OV-B5). Breeding containers located in high residential density areas with a higher flood risk, despite being watered by rainwater or water for domestic usage, might have a higher chance to be reached by flood waters. However, flood waters can also extend the range of potential habitats for mosquitoes (Yee, et al. 2019). Even more unusual breeding sites such as underground septic tanks were reported as favorable for *Ae. aegypti* reproduction in residential areas in Puerto Rico (R. Barrera, et al. 2008). These conflicting potential effects of flood on mosquito habitats might explain the higher vulnerability for mosquito occurrence in high residential density areas with higher flood risks (node OV-B6) compared with high residential density areas with lower flood risks (node OV-B5).

Node OV-B1, which had less precipitation (≤0.45 mm/hr), residential roads (≤0.02 m/m^2^), and no health institution areas (≤0%), is an extreme situation of null sensitivity of the OI toward high residential density areas. This node’s slope (0.00, R^2^ = 0.00, p = 0.84) might indicate that the mixture of other types of landscapes with residential areas is essential to enhance the sensitivity of the OI in high residential density areas. However, further work is necessary to test this hypothesis and explain potential mechanistic effects.

The adaptation of *Ae. aegypti* closer to human settlements does not seem to be solely influenced by environmental factors; certain human practices (e.g., housing in flood prone areas, weak environmental sanitation, obstruction of drainage canals, water storage practices for domestic usage) might contribute to the occurrence of mosquitoes. Therefore, environmental improvement and integrated control measures at the community level to improve the environment surrounding households, careful domestic water storage, and other sanitation practices are the most promising solutions for reducing the occurrence of mosquitoes.

The environmental conditions associated with mosquito occurrence must be carefully interpreted given the nature of the mosquito occurrence data (OI) utilized in this study. The OI is based on the percentage of positive ovitraps and can detect the presence or absence of vectors. However, it has limited capacity in displaying the precise range of mosquito density in the environment (Harburguer, et al. 2016). Therefore, the environmental conditions inferred from this ovitrap MOB tree might only display the conditions for mosquito oviposition and not necessarily the conditions influencing mosquito abundance.

### Interactive effects between precipitation and landscapes in modulating dengue incidence

Both RF (Figure 2b) and MOB tree (Figure 4a) analyses showed a significant influence of precipitation with a combinatory influence of high residential density and commercial areas in increasing dengue incidence. The significant influence of precipitation agrees with previous studies in the Philippines (Carvajal, Viacrusis, et al. 2018, Su 2008) and Malaysia (Dickin, Schuster-Wallace and Elliott 2013), which reported precipitation as a main driver of the temporal variation of dengue incidence. These studies assumed that the high correlation between precipitation and dengue incidence is due to the increasing mosquito density during the rainy season. On the MOB tree (Figure 4a), precipitation was the partitioning variable on the first and second levels whereas high residential and commercial areas were selected as partitioning variables on level 3. Overall, dengue incidence MOB trees support the significant influence of precipitation on dengue incidence.

Previous studies have discussed the impact of precipitation on the increased number of dengue cases because of its role in increasing the number of available breeding sites for mosquitoes (Carvajal, Ho, et al. 2019, Pasin, et al. 2018). However, the transference of viruses from mosquitoes to humans occurs when the dynamics between the vector, the virus, and hosts are favorable (de Melo, Scherrer and Eiras 2012). These conditions include the ratio of infected mosquitoes, human behavior, human exposure, and the ability of mosquitoes to transfer pathogens to humans (Thi, et al. 2017). Human behavior and exposure can also be altered by meteorological conditions (Akter, et al. 2017). Precipitation combined with floods, for example, can generate changes in the physical environment such as damaging shelters, which can alter urban dynamics (e.g., population displacement, increase of human exposure to vectors), thus favoring dengue transmission (Few, et al. 2004).

In the variable selection step, precipitation showed a very strong influence on dengue incidence compared with other environmental factors. We conducted further analyses to evaluate the combinatory influence of precipitation with other environmental factors in modulating dengue incidence (Figure 4b). The association between precipitation and dengue incidence was further notable in these analyses, particularly in areas covered by high residential and commercial areas, suggesting a vulnerability of these landscapes in modulating dengue incidence. The highest slope (0.44, R^2^ = 0.24, p = 0.00) was reported for node DI-B7 (Figure 4b), which incorporates interactions between precipitation, commercial areas (>4.41%), and high residential density areas (between 25.38 and 83.97%). Herewith, this nodal pathway was considered the most influential environmental condition for increasing dengue incidence with high precipitation. The second highest slope (0.39, R^2^ = 0.17, p = 0.00) was reported for node DI-B4 and formed with environmental conditions of commercial areas (0.31 < commercial areas ≤ 4.41%) and high residential density areas (>10.90%). With these influential patterns observed in terminal nodes DI-B7 and DI-B4, we suggest that certain ecological factors in commercial and high residential areas can enhance dengue transmission with precipitation. Previous studies have shown that residential and commercial areas experience the most damaged houses during extreme rainfall and flood events in Metropolitan Manila (Porio 2014, a, Porio 2011, b). Damage to families’ shelters, followed by massive displacements, might subject many people to deteriorated conditions with limited capacity to observe disease prevention and vector control measures. The high human exposure to vectors in these areas might create an avenue for high dengue transmission. A highly sensitive influence of precipitation on increasing dengue incidence was observed under high flood risk conditions (DI-B6 and DI-B2). These terminal nodes showed higher slopes of 0.41 (R^2^ = 0.16, p = 0.00) and 0.39 (R^2^ = 0.16, p = 0.00), respectively. Conversely, terminal nodes with lower flood risk (nodes DI-B1 and DI-B5, with slopes of 0.28 (R^2^ = 0.14, p = 0.00) and 0.33 (R^2^ = 0.12, p = 0.00), respectively) displayed less sensitivity in increasing dengue incidence. These examples illustrate that the vulnerability for dengue transmission in residential and commercial areas with high precipitation can be further enhanced by floods. Floods can contribute to increased mosquito density and force people to live confined in deteriorated conditions of habitability with high exposure to vectors. The combination of human presence and exposure to vectors has been linked to high dengue transmission in residential (Scott and Morrison 2010, de Moura Rodrigues, et al. 2015) and commercial areas (Honório, et al. 2009, Thammapalo, et al. 2007). Due to the anthropophilic nature of *Ae. Aegypti*, high human presence and exposure in these areas may increase feeding opportunities for mosquitoes and increase the chances of dengue fever infections (Koyadun, Butraporn and Kittayapong 2012). Because many people are exposed in areas with high precipitation levels, it becomes easier for mosquitoes to bite and infect many people in a short time, thus increasing the incidence of dengue (Akter, et al. 2017).

We expected that the resulting dengue incidence and ovitrap MOB trees (Figure 3a and 3b) would yield similar tree topology patterns. This expectation assumed that the high dengue incidence during the rainy season is a result of high mosquito abundance influenced by precipitation. However, our result was contrary to our assumption and could be explained by methodological limitations. The OI is based on the percentage of positive ovitraps with the presence or absence of vectors; however, it has a limited capacity in precisely displaying the range of mosquito density in the environment transmitting dengue. Although ovitraps are fast and cost-effective tools for monitoring the presence of mosquitoes, the OI has a low association with dengue incidence compared with adult mosquito abundance data (de Melo, Scherrer and Eiras 2012). Additional correlation analyses (data no shown) of the OI and dengue incidence from the terminal nodes of Figure 3a and 3b were not significantly correlated. Therefore, the mosquito occurrence data (OI) utilized in this study might be responsible for the dissimilarities in the ovitrap and dengue incidence MOB tree topology patterns.

### Accessibility of data, modeling approach, and limitations

Most previous epidemiological studies faced limitations when integrating climatic and landscape data given the scarcity of data and modeling techniques. Although we consider landscape to be static in this study, the model development did not distinguish dynamic or static terms. Since the physical characteristics of each village did not change significantly over the three years, we assumed that these characteristics remained the same for each month of the study period. Based on this assumption, we utilized a data structure from previous studies that repeated the values of the static variables for the monthly values of the climate variables in each village (Kaul, et al. 2018). This design allowed us to analyze dengue epidemiology as a function of the combinatory influences of climate dynamics over the static landscape.

Many methods from conventional statistics and machine learning may, in principle, be used to handle datasets with temporal and spatial dimensions. Usually a statistical model suggests empirical relationships between variables to generate specific outcomes based on certain assumptions and a priori knowledge of the modeled dynamics (Bzdok, Altman and Krzywinski 2018, Kapwata and Gebreslasie 2016). By contrast, machine learning does not require a specific model structure in advance. The algorithm itself can automatically utilize the original input data to identify hidden patterns in complex data structure (Richter and Khoshgoftaar 2018). Beforehand, statistics requires us to declare a formal model that incorporates our knowledge of the system. Thus, before implementing a model, careful inspection of the data is necessary (e.g., normal distribution) (Olden, Lawler and Poff 2008). Machine learning makes minimal assumptions about the data structure and can be effective even when the data are gathered without a carefully controlled experimental design (Bzdok, Altman and Krzywinski 2018). Additionally, machine learning is less sensitive to outliers and can efficiently address higher dimensionality variables even in the presence of complicated nonlinear interactions among covariates (Olden, Lawler and Poff 2008). The increase in data complexity may inherit some disadvantages to classical statistical methods. Instead, we utilized a machine learning approach such as RF for variable selection and recursive partitioning for subset selection because of their ability to handle complex datasets and evaluate nonlinear relationships in the data without having to satisfy the restrictive assumptions required by conventional approaches.

Machine learning, specifically RF, is often the preferred modeling method in a wide variety of epidemiological studies due to its capability to handle large datasets and accurately identify the best predictors (Kapwata and Gebreslasie 2016). However, many studies only ranked the relative importance of individual variables in influencing mosquitoes and dengue. Ranking the importance of variables alone is not enough to infer the dynamics occurring in the environment and their influence on dengue epidemiology (Sallam, et al. 2017). As discussed previously, there are multiple interacting factors in the environment that could play important roles in influencing mosquito and dengue occurrence. RF was able to handle the initial dataset and screen the most relevant variables associated with the OI and dengue incidence. Although RF can identify the most important variables, it cannot explain the interactions among covariates. Therefore, a recent study recommends also applying a machine learning method that is relatively easy to interpret (Ryo, Angelov, et al. 2020). By utilizing the recursive partitioning method in this study, we demonstrated important mechanistic interplays between environmental factors and presented specific conditions influencing the OI and dengue incidence. Furthermore, the same variables were identified as most influential on dengue and mosquitoes in RF and recursive partitioning. This consistency of results indicates the appropriateness of the adopted study design in capturing combinatory influences among environmental factors. However, since the data utilized are limited (July-December), subsequent studies should be conducted to infer whether complete data (January-December) with a similar modeling approach leads to different or similar results.

The utilization of RS in our study made it possible to access spatiotemporal temperature, precipitation, and vegetation data for each village. These types of data overcome the limited accessibility of such information at fine scales in areas where the spatial coverage of weather stations is coarse (German, et al. 2018). By using Google Earth Pro, we were able to use cloud computation to conduct all preliminary processing of the data, which significantly shortened the working time. Moreover, detailed LU maps can distinguish the risk of dengue and mosquito occurrence at a fine scale. Many studies that used coarse LU classification, for example, reported that *Ae. aegypti* and dengue incidence are positively correlated with residential areas (Vanwambeke, Lambin, et al. 2007, Vanwambeke, N.Bennett and Kapan 2011, Sarfraz, et al. 2012). In this study, we demonstrated that the distributions of mosquito and dengue can also vary depending on the type of density in these residential areas.

Our study, however, has certain limitations. As mentioned in the methods, the entomological data used in this study are incomplete and only correspond to the months of July-December of 2012– 2014. This period covers the rainy season in Metro Manila. Therefore, the lack of dry season data (January-June) causes potential bias in our study. Furthermore, ovitraps were installed in only 298 of 464 villages across Metro Manila. Complete data may better describe mechanistic understanding of associations between the dengue metrics and ambient environment factors. Nevertheless, our findings may still reflect the actual circumstances of LU and climatological characteristics of the study area. The results of the combinatory influences of landscape and climate may differ in other urban cities, particularly in rural areas in the Philippines and other dengue-endemic countries. Nonetheless, the methodology presented in this study can infer interplays between climate and landscape on mosquito occurrence and dengue incidence.

Our study demonstrated notable combinatory effects between climate and landscape in relation to the occurrence of mosquitoes and dengue incidence. It suggests discordant patterns wherein the OI is primarily influenced by landscapes and modulated by the effects of precipitation. Dengue incidence is primarily influenced by precipitation and modulated by landscapes. Such understanding and knowledge can further strengthen the design and implementation of prevention and control measures against dengue vectors and disease.

In recent years, vector control efforts in eliminating mosquito breeding sites have intensified in residential areas by identifying and destroying breeding sites. However, we demonstrated that the existence of potential breeding sites in the landscape is not the only reason for dengue transmission. These efforts should be accompanied by effective improvements in urban settings by making them more resilient to mosquito-vectored diseases.

## Acknowledgments

We thank Katherine M. Viacrusis, Lara Fides T. Hernandez, and Howell T. Ho from Trinity University of Asia for collecting and providing the data on notified dengue fever cases. This study was supported in part by the Japan Society for the Promotion of Science (JSPS) Grant-in-Aid Fund for the Promotion of Joint International Research (Fostering Joint International Research (B)) under grant number 19KK0107, JSPS Grant-in-Aid for Scientific Research (A) under grant number 19H01144, the JSPS Core-to-Core Program B. Asia-Africa Science Platforms, and the Endowed Chair Program of the Sumitomo Electric Industries Group Corporate Social Responsibility Foundation.

The authors declare they have no actual or potential competing interests.

## Data sharing

Entomological data are available from the Department of Science and Technology (http://oltrap.pchrd.dost.gov.ph/); epidemiological data are available from the National Epidemiology Center, Department of Health–Philippines, upon request; meteorological (Precipitation and Temperature) and vegetation data from remote sensing are available through GEE (https://code.earthengine.google.com/); land use data from the Geoportal Philippine website (www.geoportal.gov.ph) are available upon request; population statistics of Metro Manila are available from Philippine Statistics Authority (http://www.psa.gov.ph); and flood risk data are available from LiDAR Portal for Archiving and Distribution website (https://lipad.dream.upd.edu.ph).

## Notes

### Competing Interest Statement

The authors have declared no competing interest.

